# Identification of two novel heterodimeric ABC transporters in melanoma: ABCB5β/B6 and ABCB5β/B9

**DOI:** 10.1101/2022.10.21.513191

**Authors:** Louise Gerard, Laurent Duvivier, Marie Fourrez, Paula Salazar, Lindsay Sprimont, Di Xia, Suresh V. Ambudkar, Michael M. Gottesman, Jean-Pierre Gillet

## Abstract

ABCB5 is a member of the ABC transporter superfamily composed of 48 transporters, which have been extensively studied for their role in cancer multidrug resistance and, more recently, in tumorigenesis. ABCB5 has been identified as a marker of skin progenitor cells, melanoma, and limbal stem cells. It has also been associated with multidrug resistance in several cancers. The unique feature of ABCB5 is that it exists as both a full transporter (ABCB5FL) and a half transporter (ABCB5β). Several studies have shown that the ABCB5β homodimer does not confer multidrug resistance, in contrast to ABCB5FL. In this study, using three complementary techniques; (1) nanoluciferase-based bioluminescence resonance energy transfer, (2) co-immunoprecipitation, and (3) proximity ligation assay, we identified two novel heterodimers in melanoma: ABCB5β/B6 and ABCB5β/B9. Both heterodimers could be expressed in high-five insect cells and ATPase assays revealed that they have a basal ATPase activity. These results are an important step toward untangling the functional role of ABCB5β in melanocytes and melanoma.

## Introduction

ABCB5 is an ATP-binding cassette (ABC) transporter that is a member of one of the largest, most ancient superfamilies of proteins found in all living organisms. Mammalian ABC proteins are classified into seven families from A to G based on sequence homology(1). In humans, ABC transporters are mostly exporters, with the exception of ABCA4(2) and ABCD4(3), which are importers. They exist either as full transporters with two transmembrane domains (TMDs) and two nucleotide binding domains (NBDs) or as half transporters with one TMD and one NBD, which must either homodimerize or heterodimerize to become functional. However, not all are transporters. ABCE and ABCF members exist as twin NBDs without TMDs and are involved in mRNA translation control(4). In the ABCC family, ABCC7 is an ATP-gated chloride channel, also known as the cystic fibrosis transmembrane conductance regulator (CFTR)(5), while ABCC8 and ABCC9 serve as regulatory subunits of the ATP-sensitive potassium channel (Kir6.x)(6).

ABCB5 has been highlighted as a marker of skin progenitor cells(7), melanoma stem cells(8), and, more recently, limbal stem cells(9). It has also been reported to be a mediator of multidrug resistance in melanoma(10), breast cancer(11), colorectal cancer(12, 13), hepatocellular carcinoma(14), and several hematological malignancies(15, 16). Despite these reports, characterization of ABCB5 remains minimal. Several transcripts of human ABCB5 have been identified and transcribed from at least two different promotors, as reported in the Zenbu genome browser. According to the AceView program, which provides a strictly cDNA-supported view of the human transcriptome and the genes, ABCB5 gene transcription gives rise to at least 11 different transcript variants(17). Among these, three variants have been documented: ABCB5.b(18) (812 aa, also referred to as ABCB5β), ABCB5.h(19) (131 aa, also referred to as ABCB5α), and ABCB5.a(20) (1257 aa, also referred to as ABCB5FL Full-Length).

ABCB5β was predicted to have a TMD composed of six α-helices flanked by two intracellular NBDs(21). Conventional half transporters possess only one NBD, either at the N- or the C-terminal region(22). The ABCB5β N-terminal NBD lacks the Walker A motif, which precludes the binding of ATP. However, ABCB5β could dimerize to form a functional transporter. Potential dimerization motifs have been identified in its N-terminal region(21). Nevertheless, it has been shown that an ABCB5β homodimer cannot confer drug resistance in either the yeast model *S. cerevisiae* or in mammalian cells (20, 23). Since these studies focused on a limited number of drugs, we cannot exclude the possibility that this homodimer might be involved in drug resistance or biological functions that have yet to be unraveled. Yet, it is also worth considering that ABCB5β could dimerize with other half-transporters of the ABCB family to become functional.

Interestingly, ABCB5β belongs to a family that comprises both full and half transporters. These latter include ABCB2/TAP1, ABCB3/TAP2, ABCB6, ABCB7, ABCB8, ABCB9/TAPL, and ABCB10. The first two, ABCB2 and ABCB3, are known to dimerize in the membrane of the endoplasmic reticulum (ER) to transport peptides from the cytoplasm into the ER lumen, where they associate with the major histocompatibility complex (MHC) class I molecules and are presented to T-lymphocytes(24). Furthermore, ABCB7 and ABCB10 have been shown to form a complex with ferrochelastase in the mitochondria, where they are involved in heme biosynthesis(25).

The three remaining half transporters have been linked to multidrug resistance in several cancer types, including melanoma(26-28). Bergam and colleagues demonstrated that ABCB6 is involved in the early steps of melanogenesis(29). Furthermore, mutations in this gene are responsible for dyschromatosis universalis hereditaria, a rare pigmentary genodermatosis(30). ABCB5 and ABCB8 have been proposed to play a role in the transport of melanin metabolites(31). These transporters could mediate the sequestration of toxic melanin metabolites into lysosomes or melanosomes, thereby protecting melanocytes. In melanoma, they could sequester drugs in these organelles(32). To date, no strong experimental evidence supports these conjectures. Considering that several ABCB half transporters form functional heterodimers, we hypothesized that ABCB5β might heterodimerize with other related members of the ABCB family, possibly, ABCB6, ABCB8 or ABCB9.

In this study, we identified two novel heterodimeric ABC transporters in melanoma, ABCB5β/B6 and ABCB5β/B9, using three complementary techniques: nanoluciferase-based bioluminescence resonance energy transfer (NanoBRET), co-immunoprecipitation, and proximity ligation assay (PLA). When ABCB5β/B6 and ABCB5β/B9 heterodimers were fused with P-glycoprotein flexible linker region and expressed in high-five insect cells, they exhibited significant levels of basal ATPase activity. This discovery paves the way for a better understanding of the roles played by these heterodimeric transporters in melanoma.

## Results

### NanoBRET assay reveals ABCB6 and ABCB9 as interacting partners of ABCB5β

We hypothesized that ABCB5β might heterodimerize with a half transporter of the ABCB family. The review of the literature, which shows ABCB2-ABCB3 and ABCB7-ABCB10 interaction, led us to select three half transporters, namely ABCB6, ABCB8, and ABCB9, as possible partners for ABCB5β. To study the putative interactions between ABCB5β and these proteins, we conducted a NanoBRET assay as described in **Supporting information Fig. S1**. Each of the four proteins was tagged with either a NanoLuc-donor or HaloTag-acceptor at either the amino (N) or carboxy (C) terminus of the protein, resulting in 16 constructs and eight potential donor/acceptor combinations for each protein pair **(Supporting information Fig. S1B)**. For each combination to be tested, two constructs (one with NanoLuc and one with HaloTag) were transfected in HEK-293T cells, and either the fluorescent NanoBRET HaloTag 618 Ligand or a DMSO vehicle was added to the cells. The NanoBRET NanoLuc substrate was then added, and donor and acceptor signals were measured, which allowed us to calculate a NanoBRET ratio by dividing the acceptor emission value by the donor emission value for each sample in both conditions. To factor in the donor-contributed background, we subtracted the no-ligand control NanoBRET ratio (i.e., DMSO vehicle) from the ligand-containing sample NanoBRET ratio to obtain the corrected NanoBRET ratio **(Supporting information Fig. S1B)**.

As controls for the assay, we also tested the interaction between additional ABC proteins. To this end, we tested the fully characterized ABCB2/B3 heterodimer as a positive control(24). Furthermore, we tested the putative interactions between ABCB2-ABCD1 and ABCB3-ABCD1 proteins as negative controls, since the ABCB2/B3 heterodimer is expressed in the ER membrane while ABCD1 is a peroxisomal transporter. As an additional control, we also looked for a possible interaction between ABCB5β and ABCD1 proteins.

Each tested protein pair yielded a high NanoBRET ratio for one or more donor/acceptor combinations, suggesting heterodimerization (**Supporting information Fig. S1B**). Variations in the transfection efficiency of the donor and acceptor constructs or the constitutive expression of the proteins lead to an environment where the concentrations of all partners were not regulated, rendering any quantitative assumption difficult. As a result, in this experiment, the NanoBRET ratio was used as qualitative information. Ratios similar to the no-ligand negative control were classified as noninteracting, while ratios above that suggested heterodimerization.

It should be noted that a major drawback of resonance energy transfer assays (e.g., FRET, BRET, and NanoBRET) is that a signal can be detected when proteins are in close proximity but do not interact together(33). This is typically the case for transmembrane proteins, as under overexpression conditions, membrane crowding may lead to protein aggregates where donors and acceptors can transfer energy to one another without the generation of heterodimers. To overcome this limitation, we decided to decrease the amount of donor and acceptor constructs linearly. Nonspecific interaction should result in decreased NanoBRET ratios, while these ratios should remain stable if the interaction is specific.

For this experiment, we chose the best donor/acceptor combination based on the highest NanoBRET ratio (**Supporting information Fig. S1B**). The absence or low signal in alternative tag combinations for a tested pair indicates that the tags might be either too far from one another to transfer energy, not correctly oriented, or might be impairing protein folding, localization, or dimerization(34). Therefore, the pair with the largest NanoBRET ratio represents the combination for which both tags are in the best conformation to transfer energy to one another. The selected pairs are highlighted in **Supporting information Fig. S1B**.

HEK-293T cells were transfected with decreasing amounts of NanoLuc (donor) and HaloTag (acceptor) constructs ranging from 1 µg to 0.125 µg for each construct (**Fig. 1**). The NanoBRET ratio decreased, along with the DNA amount, for ABCB2-ABCD1, ABCB3-ABCD1, and ABCB5β-ABCD1 pairs tested as negative controls (**Fig. 1A**) and stayed constant regardless of the amount of DNA transfected for ABCB2-ABCB3 used as a positive control (**Fig. 1B**). Interestingly, NanoBRET ratios remained similar in all conditions for ABCB5β/B6 and ABCB5β/B9, suggesting that these proteins might form true heterodimers. The NanoBRET ratio obtained for ABCB5β-ABCB8 decreased along with the amount of DNA transfected, similar to the data generated from protein pairs assessed as negative controls.

**Fig. 1:**
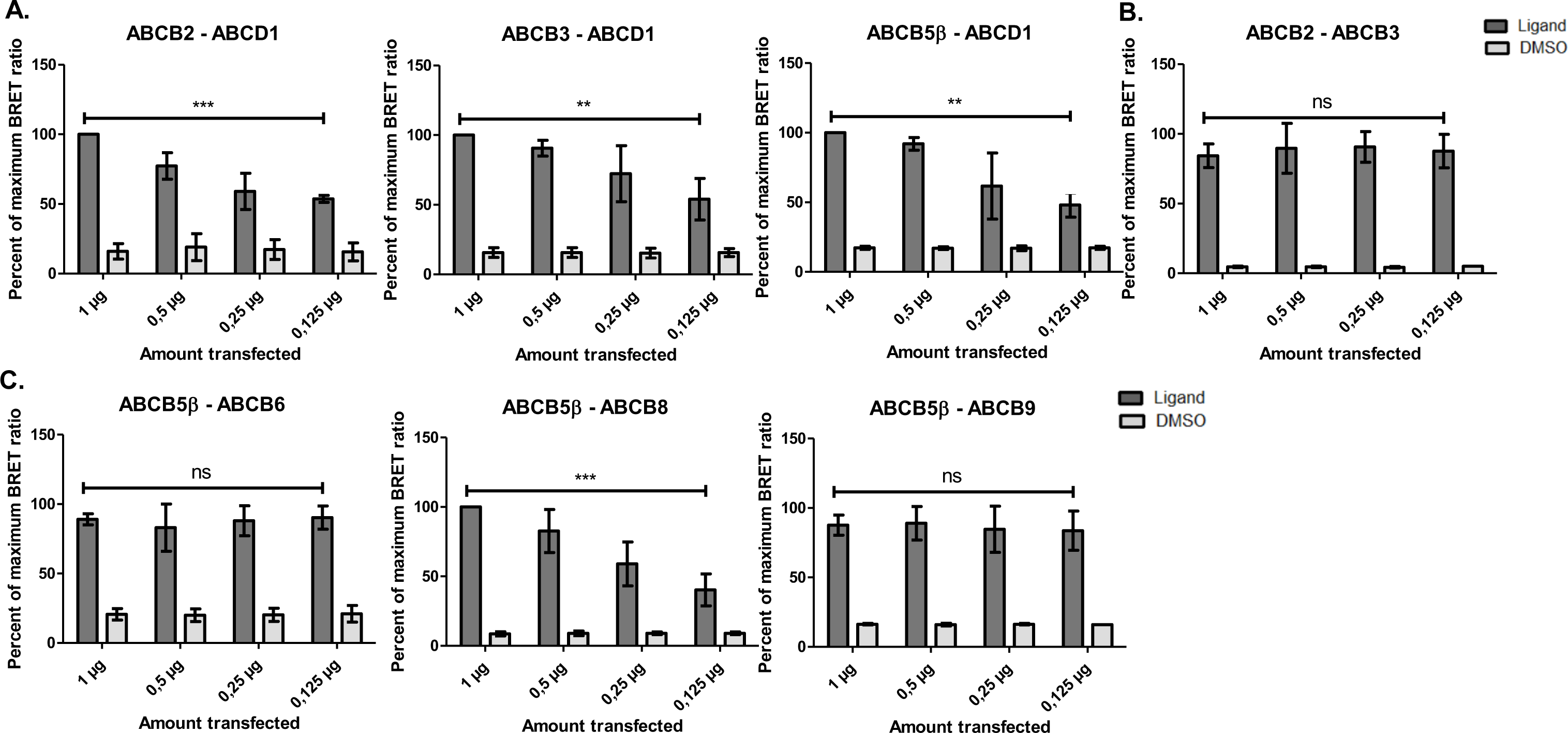
NanoBRET dilution assay reveals two putative heterodimers, ABCB5β/B6 and ABCB5β/B9. HEK-293T cells were transfected with decreasing amounts of NanoLuc and HaloTag constructs. Transfection started at 1 µg and gradually decreased to 0.125 μg per construct. DMSO refers to the technical negative control to which DMSO is added instead of ligand. Protein pairs serving as negative controls (ABCB2-ABCD1, ABCB3-ABCD1, ABCD1-ABCB5β) (**A**) and a positive control pair (ABCB2-ABCB3) (**B**) were tested alongside ABCB5β-ABCB6, ABCB5β-ABCB8, and ABCB5β-ABCB9 (**C**). Data are presented as mean ± SD. Simple linear regression was used to test whether the amount transfected significantly influenced the NanoBRET ratio (*n* = 3). *P*-values are represented on graphs as *** *p* < 0.001, ** *p* < 0.01, and *p* > 0.05. For ABCB2-ABCD1 (*R*^2^ = 0.8719, *F*(1,10) = 68.07), ABCB3-ABCD1 (*R*^2^ = 0.6144, *F*(1,10) = 15.94), ABCB5β-ABCD1 (*R*^2^ = 0.6661, *F*(1,10) = 19.95), and ABCB5β-ABCB8 (*R*^2^ = 0.7526, *F*(1,10) = 30.41), the relation between both parameters was statistically significant. For ABCB2-ABCB3 (*R*^2^ = 0.0269, *F*(1,10) = 0.2767), ABCB5β-ABCB6 (*R*^2^ = 0.0011, *F*(1,10) = 0.0113), and ABCB5β-ABCB9 (*R*^2^ = 0.0204, *F*(1,10) = 0.2080), changes in the amount of transfected DNA did not influence the NanoBRET ratio.

### Donor saturation assay confirms the specificity of the interaction between ABCB5β/B6 and ABCB5β/B9 protein pairs

To confirm the specific interaction of the ABCB5β-ABCB6 and ABCB5β-ABCB9 protein pairs and the false positive signals of the ABCB5β-ABCB8, ABCB2-ABCD1, ABCB3-ABCD1, and ABCB5β-ABCD1 protein pairs, a saturation assay was performed in HEK-293T cells (**Fig. 2**). In this experiment, the amount of donor construct was held constant, while the amount of acceptor construct was gradually increased. For this assay, the smallest concentrations of donor and acceptor showing heterodimerization were used. If the interaction between both proteins is specific, the NanoBRET signal will increase in a hyperbolic manner, reaching a plateau when all the donors are saturated with the acceptors and luminescence can no longer increase. On the other hand, for a nonspecific interaction, the signal will increase linearly with an increasing acceptor concentration (**Fig. 2A**). At these concentrations, the signal from a nonspecific interaction will not reach a plateau due to the lack of affinity between proteins.

**Fig. 2:**
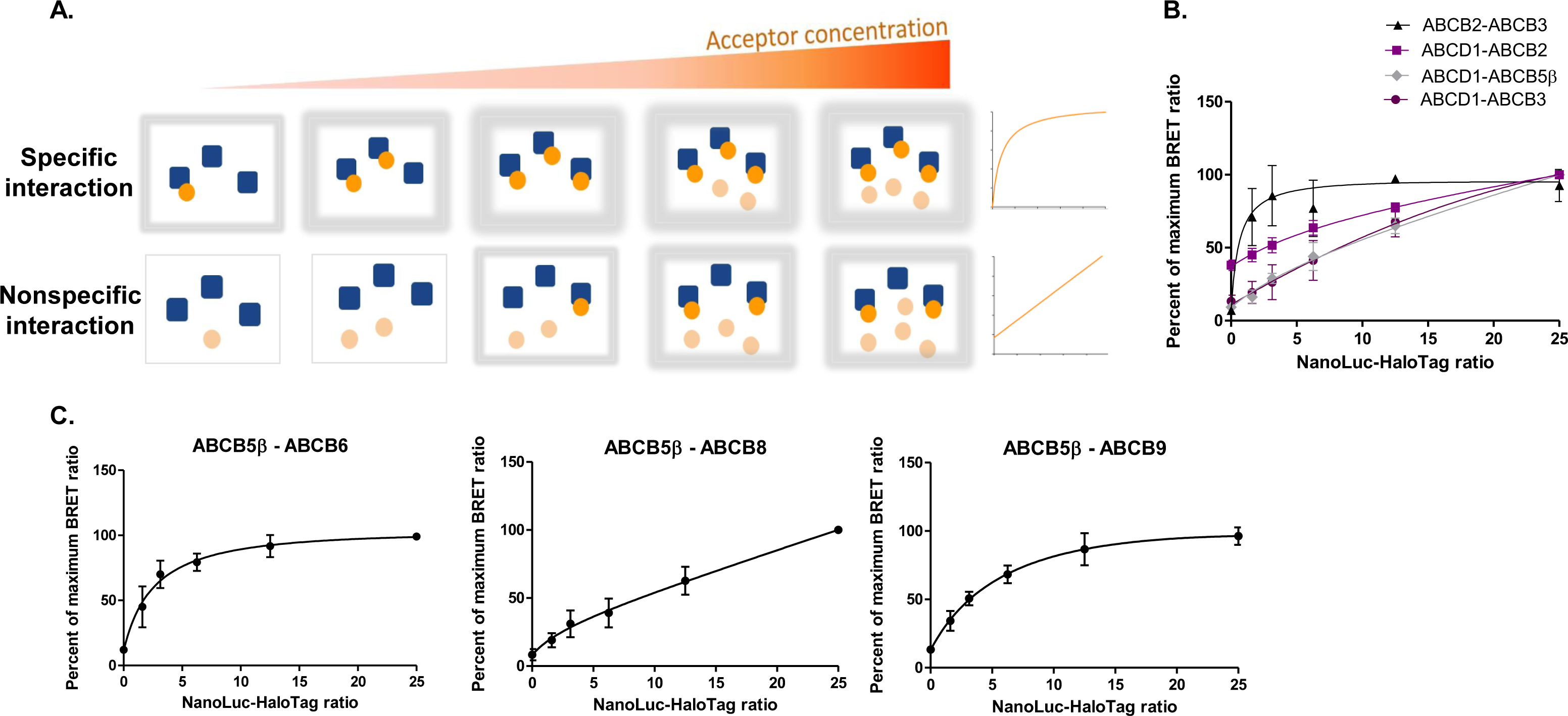
Saturation assay confirms specific ABCB5β-ABCB6 and ABCB5β-ABCB9 interactions. **(A)** The concentration of the donor is held constant while the concentration of the acceptor is increased. If the interaction is specific, the signal will reach a plateau where all donors are saturated with acceptors, and luminescence will no longer increase. In the case of a nonspecific interaction, the signal will continue to increase linearly. For each square, the schematic gray halo represents the intensity of the NanoBRET signal observed from low to high. In graphs (**B-C**), the percent of the maximum NanoBRET ratio is plotted against the NanoLuc−HaloTag ratio. (**B)** Donor saturation assay for the positive control, ABCB2-ABCB3, and the negative controls, ABCB2-ABCD1, ABCB3-ABCD1, and ABCB5β-ABCD1. (**C)** Donor saturation assay for ABCB5β-ABCB6, ABCB5β-ABCB8, and ABCB5β-ABCB9.

We carried out this donor saturation assay on the ABCB2-ABCB3 protein pair used as a positive control and compared the data to that from the three protein pairs chosen as negative controls. As shown in **Fig. 2B**, the NanoBRET signal was saturable for the ABCB2-ABCB3 protein pair while a linear increase was observed for the ABCB2-ABCD1, ABCB3-ABCD1, and ABCB5β-ABCD1 pairs. This supports the specificity of the interaction for the ABCB2-ABCB3 protein pair. The specificity of the interaction was then assessed for the three protein pairs of interest. The analysis of the data revealed a hyperbolic increase in the NanoBRET signal that eventually reached a plateau phase for both ABCB5β-ABCB6 and ABCB5β-ABCB9 protein pairs, similar to what was observed for the positive control. On the contrary, the donor saturation assay further supported a nonspecific interaction between the ABCB5β and ABCB8 proteins (**Fig. 2C**).

### Co-immunoprecipitation and proximity ligation assay confirm interactions in cells between the ABCB5β-ABCB6 and ABCB5β-ABCB9 protein pairs

Co-immunoprecipitation (Co-IP) is considered one of the standard methods for identifying or confirming the occurrence of protein-protein interaction events. Because of this, as a first step, we used HEK-293T cells which constitutively express ABCB6 and ABCB9. Since the expression of ABCB5β had not been demonstrated in this cell line, we transfected them with mCherry-tagged ABCB5β (**Supporting information Fig. S2**). The proteins of interest were immunoprecipitated using either an anti-mCherry, ABCB5, ABCB6, or ABCB9 antibody, and the interacting partners were analyzed using Western blotting after SDS-PAGE using corresponding antibodies to show protein−protein interaction (**Supporting information Fig. S2**). An isotype control was used to determine the specificity of the signal obtained in the Western blots.

Membrane crowding, protein mislocalization, misfolding, and loss of function are some of the issues that might arise from membrane protein overexpression and the utilization of a tag. Therefore, we decided to work with Mel JuSo and UACC-257, two melanoma cell lines that constitutively express the target proteins (**Fig. 3**). Immunoprecipitation with either an anti-ABCB5, ABCB6, or ABCB9 antibody detects the interacting partners, which were identified by Western blotting after an SDS-PAGE using the corresponding antibody. The data further supports the occurrence of interaction between ABCB5β-ABCB6 and ABCB5β-ABCB9 protein pairs (**Fig. 3**).

**Fig. 3:**
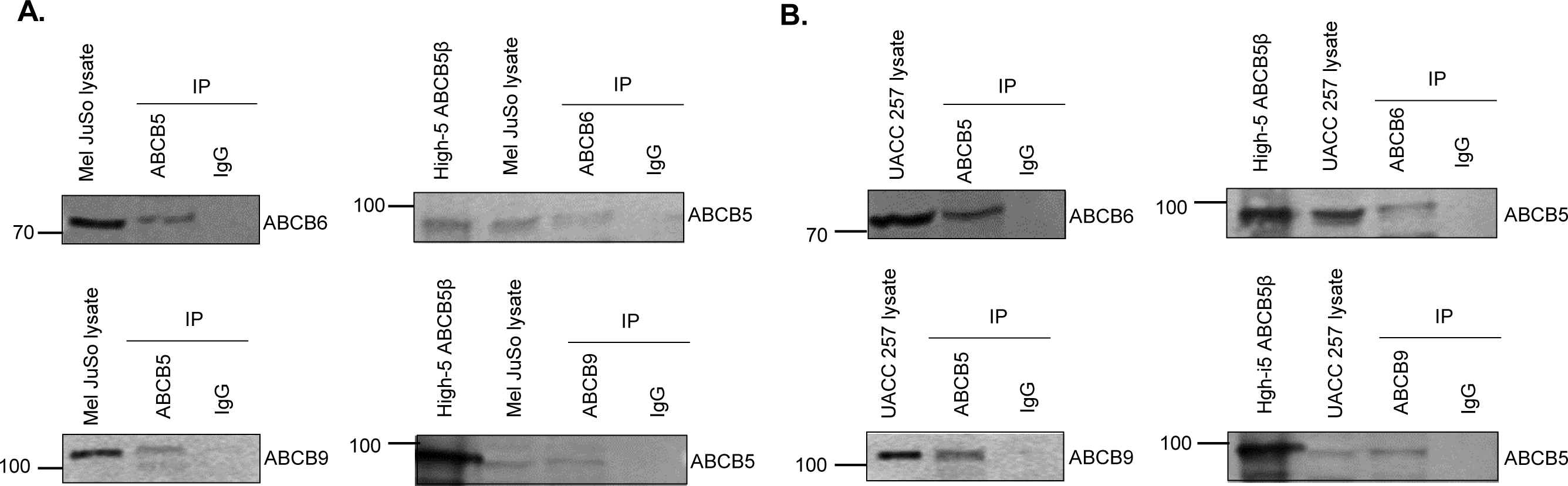
Co-immunoprecipitation of ABCB5β-ABCB6 and ABCB5β-ABCB9 demonstrates the presence of these heterodimers in Mel JuSo and UACC-257 cells. The Mel JuSo (A) or UACC-257 (B) proteins were immunoprecipitated (IP) with either an anti-ABCB5, anti-ABCB6, or anti-ABCB9 antibody. The precipitated proteins were revealed by Western blotting after SDS-PAGE using the corresponding antibody indicated on the right side of each blot. Fifteen µg of total proteins from the starting cell lysate were loaded in the first lane while the total IP eluate was loaded on the gel. ABCB5β was expressed in High Five insect cells, and total membrane proteins were prepared and loaded on the gel (High5 ABCB5β) as a complementary molecular weight marker when using anti-ABCB5. An isotype control was performed to determine the specificity of the signal obtained in Western blot.

*In situ* validation of the interactions for these two protein pairs was obtained using a proximity ligation assay (PLA) in both Mel JuSo and UACC-257. The PLA principle is illustrated in **Supporting information Fig. S3**. Strong red PLA signals were detected for ABCB5β-ABCB6 and ABCB5β-ABCB9, indicating interactions between the studied protein pairs (**Fig. 4A**). To further assess the specificity of the PLA signal, multiple controls were performed. Biological controls were conducted in which Mel JuSo and UACC-257 cell lines were produced that stably expressed either a nontargeting short hairpin RNA (shRNA) or an ABCB6-specific (or ABCB9-specific) shRNA, resulting in the decrease of either ABCB6 or ABCB9 expression. **Fig. 4B and C** show the resulting decrease of expression at the mRNA and protein levels. In both cell lines, stable knockdown of either ABCB6 or ABCB9 resulted in a statistically significant decrease in PLA signal (*p*<0.0001) as compared to the nontargeting shRNA samples (**Fig. 4D**). Technical controls were also conducted in which just one of the two antibodies directed against the proteins of interest was added, or neither was added (**Fig. 4D**).

**Fig. 4:**
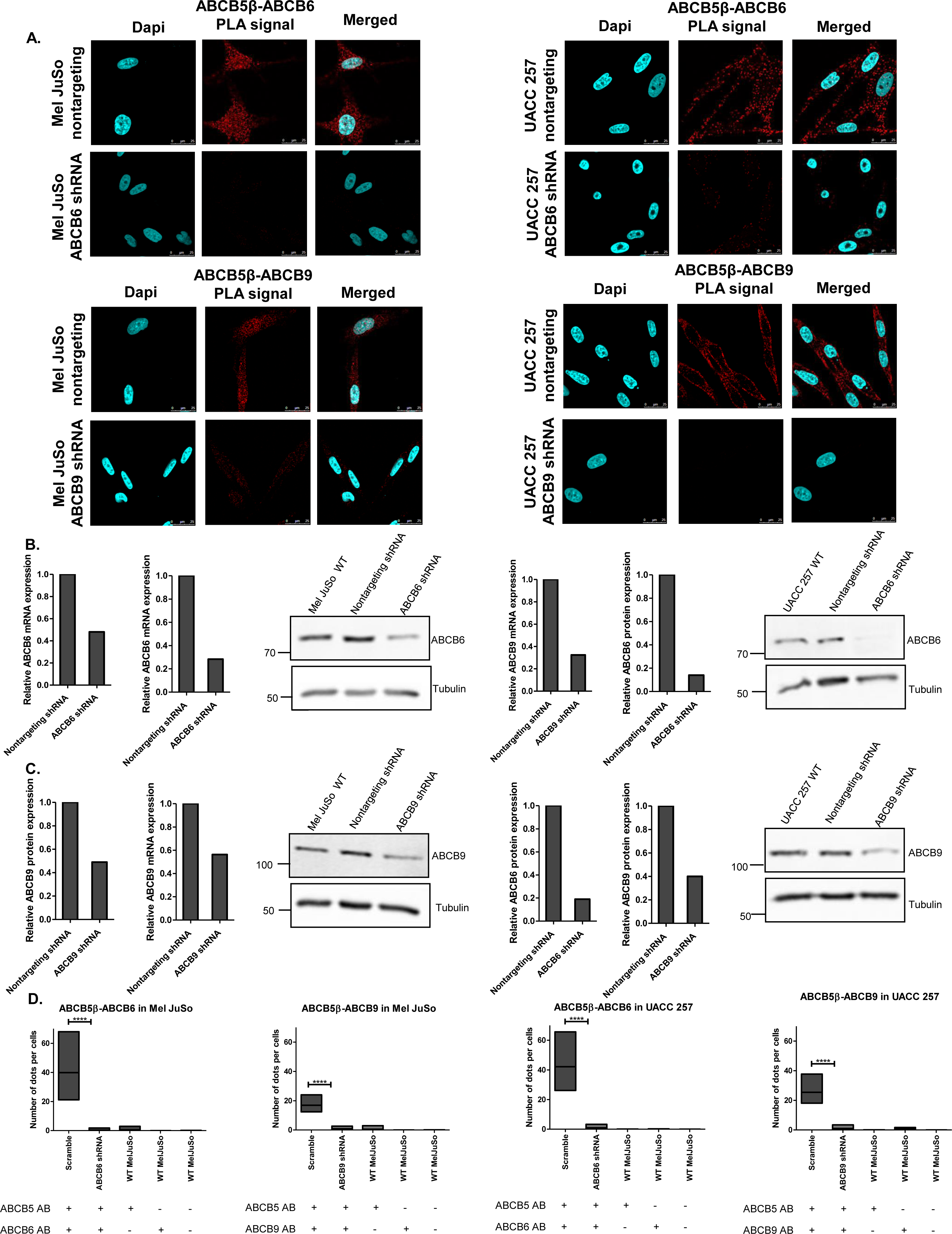
Proximity ligation assay confirms the interactions between ABCB5β-ABCB6 and ABCB5β-ABCB9 protein pairs in Mel JuSo and UACC-257. (A) Confocal images of PLA in Mel JuSo and UACC-257 that stably expressed either a nontargeting short hairpin RNA (shRNA) or an ABCB6-specific (or ABCB9-specific) shRNA. The detected PLA signal is in red. Nuclei are stained in blue using DAPI. After shRNA knockdown, efficiency was determined using RT-qPCR and Western blots in Mel JuSo (B) and UACC-257 (C). 18S and tubulin were used as references for RT-qPCR and Western blots normalization, respectively (n = 2). ImageJ was used to quantify protein expression in the Western blot as well as the average number of dots per cell (*n* = 3) (D). Scramble and shRNA quantifications are shown along with technical negative controls, including PLA conducted using either one antibody (AB) or none. Data is presented as mean ± *SD*. In Mel JuSo: ABCB6 scramble (39.89 ± 14.21), ABCB6 shRNA (0.28 ± 0.60), ABCB5 AB control (0.55 ± 0.91), ABCB6 AB control (0 ± 0), ABCB9 scramble (16.89 ± 4.27), ABCB9 shRNA (0.84 ± 1.04), ABCB9 AB control (0 ± 0.01), and no antibody control (0.01 ± 0.05). In UACC-257: ABCB6 scramble (42.16 ± 10.62), ABCB6 shRNA (0.97 ± 1.00), ABCB5 AB control (0.01 ± 0.02), ABCB6 AB control (0.03 ± 0.07), ABCB9 scramble (25.39 ± 6.74), ABCB9 shRNA (1.04 ± 1.16), ABCB9 AB control (0.79 ± 0.58), and no antibody control (0 ± 0). Student’s *t*-test was performed between the scramble and shRNA conditions. Statistical significances are presented on graph as \*\*\*\**p* < 0.0001. ABCB5β/B6 in Mel JuSo (*t* = 9.648, *df* = 22), ABCB5β/B9 in Mel JuSo (*t* = 12,66, *df* = 22), ABCB5β/B6 in UACC-257 (*t* = 13,38, *df* = 22), ABCB5β/B9 in UACC-257 (*t* = 12,34, *df* = 22).

### ABCB5β/B6 and ABCB5β/B9 linked with the P-gp linker exhibit basal ATPase activity

Next, we wanted to determine whether ABCB5β-ABCB6 and ABCB5β-ABCB9 could hydrolyze ATP, which is required for the transport of substrates across membranes. Genetic constructs were generated to ensure that we were studying the function of these transporters after heterodimerization and not homodimerization. These constructs were composed of each half transporter cDNA fused with its interacting partner using the flexible linker region of P-glycoprotein (P-gp), also known as ABCB1, comprised of 57 amino acids (residues 703N-760D). As described by Bathia *et al*., this design guarantees heterodimerization but has no effect on either the localization or behavior of the transporter (35). Moreover, due to the highly flexible nature of the P-gp linker region, its structure remains elusive as it has not been solved in any of atomic structures of this transporter. To assess all possible orientations, four different constructs were prepared (ABCB5β_P-gp linker_ABCB6, ABCB6_P-gp linker_ABCB5β, ABCB5β_P-gp linker_ABCB9, ABCB9_P-gp linker_ABCB5β). High-five insect cells were transduced using baculovirus, encoding the aforementioned recombinant DNA, and harvested 58-72 hours after infection. Total membrane vesicles were prepared as described in the methods section. To confirm the expression of the heterodimer chimeras in the insect cells, SDS-PAGE followed by Coomassie-blue staining (**Fig.5A**) or SDS-PAGE and Western blotting (**Fig.5B**) using anti-ABCB5, anti-ABCB1, anti-ABCB6, and anti-ABCB9 antibodies, were performed. Insect cells membrane vesicles expressing only ABCB5β, ABCB6, and ABCB9 were used as a control for the homodimeric form.

**Fig. 5:**
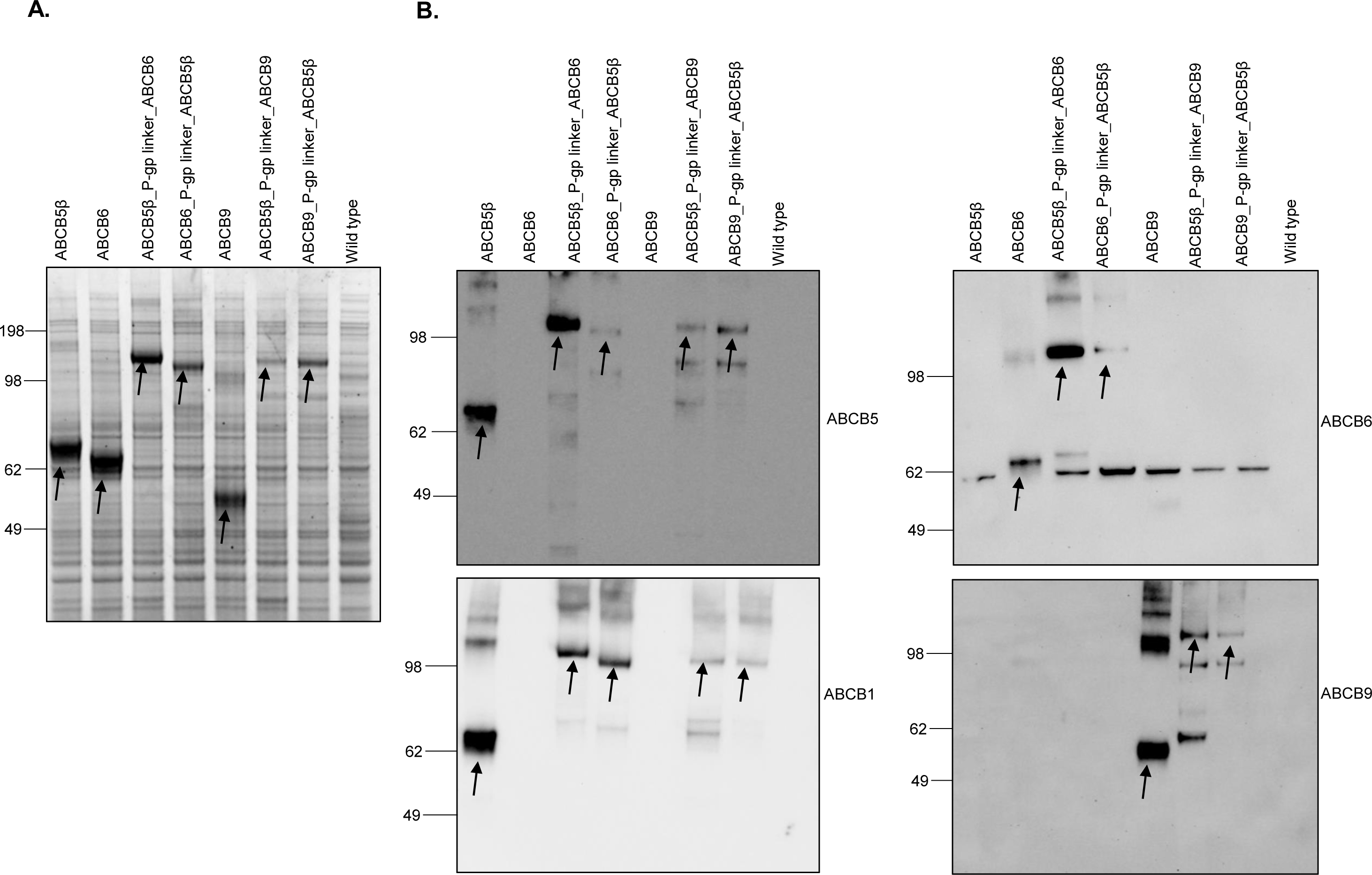
Expression of ABCB5β, ABCB6, ABCB5β_P-gp linker_ABCB6, ABCB6_P-gp linker_ABCB5β, ABCB9, ABCB5β_P-gp linker_ABCB9, and ABCB9_P-gp linker_ABCB5β in High-five insect cells. Total membranes vesicles prepared from high-five cells infected with baculovirus containing either ABCB5β, ABCB6, ABCB5β_P-gp linker_ABCB6, ABCB6_P-gp linker_ABCB5β, ABCB9, ABCB5β_P-gp linker_ABCB9, or ABCB9_P-gp linker_ABCB5β were subjected to SDS-PAGE followed by Coomassie-blue staining (A) or Western blotting with anti-ABCB5, anti-ABCB1, anti-ABCB6, and anti-ABCB9 antibodies (B). Bands of interest are highlighted by black arrows.

Keniya and colleagues demonstrated the cross reactivity of anti-ABCB1 (mouse monoclonal C219) with ABCB5β(23). **Fig. 5B** shows that anti-ABCB1 was also able to recognize ABCB5β when expressed alone or fused with its interacting partners using the P-gp linker. This is due to the significant sequence homology between the NBDs of ABCB5β and ABCB1(23). Nevertheless, ABCB6 and ABCB9 did not cross react with the anti-ABCB1 antibody. This demonstrates that the epitope of the C219 antibody is not present in the NBDs of ABCB6 and ABCB9.

The quantification of Coomassie-blue staining intensity of the monomer, ABCB5β, ABCB6, and ABCB9 showed that these transporters are expressed to similar levels in high-five insect cells (**Fig. 6A and B**). The different antibodies used, and their different reactivities, could explain the differences observed in the intensity of bands in Western blots (**Fig. 6C**). ATP hydrolysis was measured in the high-five total membrane vesicles that expressed the homodimers, by the quantification of the inorganic phosphate released in a colorimetric assay. The three homodimers showed significant ATPase activity, which was inhibited by beryllium fluoride (BeFx) (**Fig. 6D**). The three transporters showed comparable ATPase activity as well as inhibition of this activity by BeFx.

**Fig. 6:**
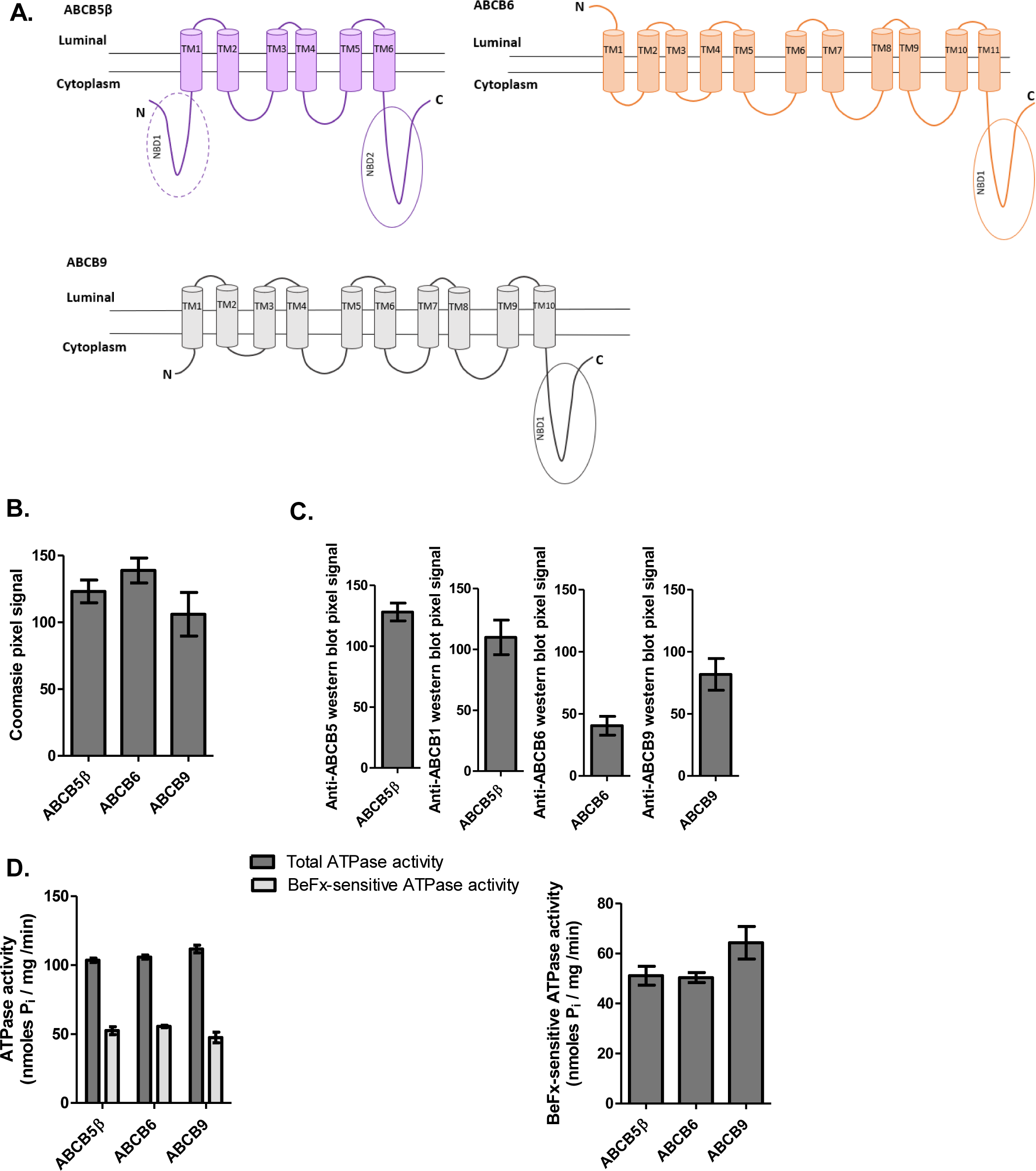
Expression of homodimers in high-five cells and BeFx-sensitive ATPase activity. (**A**) Two-dimensional schematic representation of ABCB5β (purple), ABCB6 (orange), and ABCB9 (grey) structure based on CCTOP predictions(36). ABCB5β has one complete and another partial NBD and one TMD comprised of six transmembrane helices (TM). The N terminal NBD (NBD1) is truncated and lacks the conserved Walker A domain. ABCB6 has 6 transmembrane helices (TM6-TM11) that constitutes its TMD1 and 5 additional transmembrane helices (TM1-TM5) in the N terminus that form the TMD0. ABCB9 has a TMD1 consisting of 6 transmembrane helices (TM5-10) and a TMD0 made of 4 transmembrane helices at the N terminus. Pixel intensity quantification of the protein bands of interest, after Coomassie-blue staining (**B**) or Western blotting with anti-ABCB5, anti-ABCB6, and anti-ABCB9 antibodies (**C**) was performed using ImageJ (n=3). (**D**) ABCB5β, ABCB6, and ABCB9 ATPase activities were measured in the presence and absence of BeFx as described in the method section. On the left, data are shown as ATPase activity in the absence of BeFx (total ATPase activity) and presence of BeFx (BeFx-sensitive ATPase activity). On the right, data are represented by subtracting BeFx-resistant ATPase activity from the total ATPase activity (n=3).

On the other hand, the quantification of Coomassie-blue staining and Western blot signals revealed a difference in the heterodimer’s expression levels, ranging from highest to lowest as follows: ABCB5β_P-gp linker_ABCB6, ABCB6_P-gp linker_ABCB5β, ABCB9_P-gp linker_ABCB5β, and finally ABCB5β_P-gp linker_ABCB9 (**Fig. 7A, B and C**). It is important to note that the N terminus of ABCB6 is located on the lumen side of the membrane vesicles whereas the C terminus of ABCB5β is in the cytoplasm. As a result, the CCTOP two-dimensional model assumes that the fusion of ABCB5β when at the N terminus with ABCB6 in the C terminus leads to the trapping of one transmembrane helix from ABCB6 (TM1) in the cytoplasmic space because the P-gp linker hydrophilicity prevents it from crossing the membrane. This causes ABCB6 to lose one transmembrane helix (TM1) from its TMD0 (**Fig. 7A**). Nevertheless, this conformation was expressed at a high level in high-five insect cells as compared to the three other heterodimer constructs. Interestingly, the measured ATPase activity was similar across the four heterodimeric chimeras (**Fig. 7D**).

**Fig. 7:**
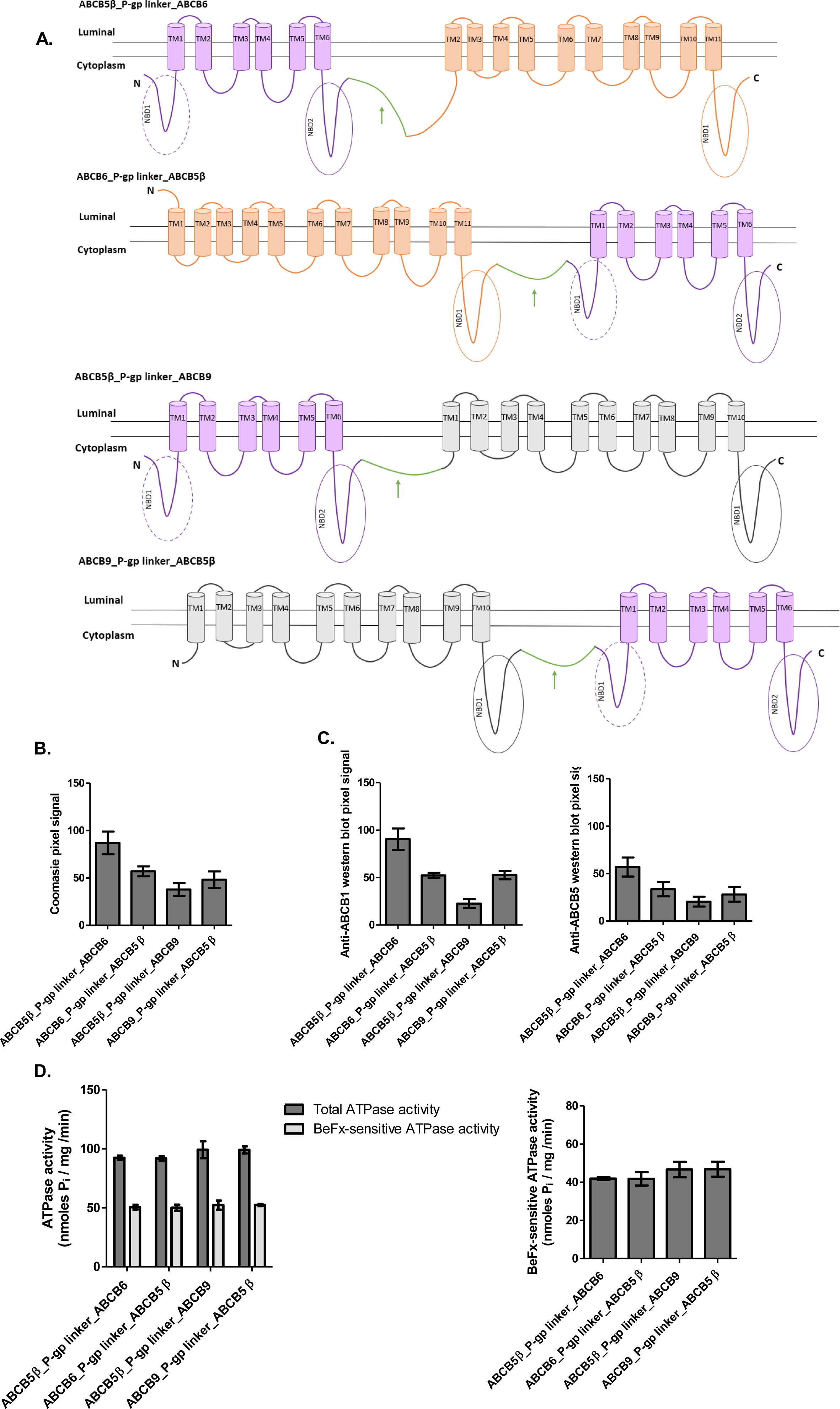
Expression level and BeFx-sensitive ATPase activity of heterodimers. (**A**) Two-dimensional schematic representation of the ABCB5β/B6 and ABCB5β/B9 heterodimers after fusion with the P-gp flexible linker based on CCTOP predictions(36). P-gp linker is shown in green and highlighted with a green arrow; ABCB5β is represented in purple, ABCB6 in orange, and ABCB9 in grey. ABCB5β_P-gp linker_ABCB6 has 16 transmembrane helices. ABCB6_P-gp linker_ABCB5β has 17 transmembrane helices. ABCB5β_P-gp linker_ABCB9 and ABCB9_P-gp linker_ABCB5β both have 16 transmembrane helices. The Coomassie-blue stained gel (**B**) or Western blotting with anti-ABCB5 and anti-ABCB1 antibodies (**C**), pixel intensity of the bands of interest were quantified using ImageJ (n=3). (**D**) The graph on the left shows the ATPase activity for all heterodimers chimeras measured in both the absence and presence of BeFx as described in the method section, total ATPase activity and BeFx-sensitive ATPase activity, respectively. On the right, data represent BeFx-sensitive ATPase activity, which is calculated by subtracting BeFx ATPase activity from the total ATPase activity (n=3).

Next, ATPase activity for all the constructs was assessed in the presence of vanadate (Vi) instead of BeFx. Lower inhibition of ATPase activity was observed with 0.3 mM Vi, resulting in a lower Vi-sensitive ATPase activity than with BeFx (**Supporting information Fig. S4**).

To visualize potential interactions, ABCB5β may have associating with itself or with others, three-dimensional models of ABCB5β homodimer and heterodimers were constructed using the approach of homology modeling, given the availability of the crystal structure of mouse ABCB1 (PDB: 5KPI) and the high sequence identity between human ABCB5β and mouse ABCB1 which is greater than 60% **(Supporting information Fig. S5**).

The significant sequence identities indicate a homologous structure and function among these transporters. The three-dimensional models indicate that the essential structural features of ABCB5β homodimer and ABCB5β/B6 and ABCB5β/B9 heterodimer are homologous with those of mouse P-gp. These models also show that the TMD0s of ABCB6 and ABCB9 are not engaged in the interaction of ABCB5β with either ABCB6 or ABCB9 **(Supporting information Fig. S5B and C)**. Importantly, these models provide a key structural framework for further research required to determine which residues of the TMDs from ABCB5β, ABCB6, and ABCB9 are involved in the interaction.

## Discussion

To explore the possible role of the ABCB5β variant, we conducted heterodimerization studies with potential target half transporters of the ABCB family in melanoma. This ABC family comprises three full-length transporters (i.e., ABCB1, B4, and B11) and seven half transporters. ABCB5 is the only member of the ABCB family that is found as both a full transporter (ABCB5FL) and a half transporter (ABCB5β), with expression driven by independent promoters. Current knowledge of these transporters led us to focus on three candidate heterodimers: ABCB5β-ABCB6, ABCB5β-ABCB8, and ABCB5β-ABCB9. Putative interactions between these three pairs of proteins were studied using the NanoBRET assay and confirmed using a co-immunoprecipitation and proximity ligation assay. Taken together, these assays revealed two novel heterodimeric ABC transporters in melanoma cell lines.

FRET microscopy in living cells has been applied to unravel the interactions of peroxisomal ABC transporters ABCD1 and ABCD3(37). In this study, we carried out a NanoBRET assay, which does not require extrinsic excitation and provides substantially increased detection sensitivity and dynamic range as compared to current BRET technologies(34). We demonstrated the importance of transfecting a low amount of donor and acceptor constructs to avoid either membrane crowding or protein aggregates that may lead to false positive BRET signals, as observed for ABCB8-ABCB5β, ABCB2-ABCD1, ABCB3-ABCD1, and ABCB5β-ABCD1 in Supporting Figure S1B. This so-called bystander signal is the result of two proteins that do not interact but are close enough to transfer energy(33). The donor saturation assay confirmed the specificity of the interaction between the ABCB5β/B6 and ABCB5β/B9 protein pairs and demonstrated the nonspecificity of the interactions for the other pairs of proteins (ABCB5β/B8, ABCB2/D1, ABCB3/D1, and ABCB5β/D1). Co-immunoprecipitation was performed to confirm the occurrence of protein-protein interaction events using the particularly easy-to-transfect HEK-293T. Since ABCB5β expression had not been observed in these cells, we transfected them with mCherry-tagged ABCB5β. This experiment allowed us to confirm the data obtained from the NanoBRET assay using the same cell line. We wanted to assess protein–protein interaction in a relevant model with a native expression of the three transporters of interest in order to avoid any potential changes in protein localization and membrane crowding caused by protein tags and protein overexpression. Since ABCB5β is predominantly expressed in melanoma, we used two melanoma cell lines, Mel JuSo and UACC-257, which also constitutively express ABCB6 and ABCB9. The goal was to validate the data *in situ* in live melanoma cell lines. As such, we carried out a proximity ligation assay in melanoma cell lines that stably express either a nontargeting short hairpin RNA (shRNA) or an ABCB6-specific (or ABCB9-specific) shRNA. This is the only experiment that provides qualitative and quantitative information. The quantification of individual PLA signals, represented by red dots, revealed that ABCB5β/B6 is more abundant than ABCB5β/B9 in both tested cell lines. Altogether, this set of experiments demonstrates the existence of the two heterodimeric ABC transporters in the Mel JuSo and UACC-257 melanoma cell lines.

To study ABCB5β heterodimers, four constructs were generated by fusing each of the interacting transporters with the P-gp flexible linker region in one of the two possible orientations, ABCB5β in either N or C terminus. After expression in high-five insect cells, each heterodimer expression was assessed and quantified using Coomassie-blue staining and Western blotting. It appears that the level of expression is dependent on the orientation for each heterodimer (ABCB5β/B6 or ABCB5β/B9). The ABCB5β_P-gp linker_ABCB6 and ABCB9_P-gp linker_ABCB5β heterodimers are expressed at higher level than others (see Fig. 7). It is important to note that, as highlighted by the CCTOP two-dimensional model, the orientation in ABCB5β_P-gp linker_ABCB6 leads to the loss of one transmembrane helix from ABCB6 TMD0, which becomes an intracytoplasmic loop (or α-helical region). However, the transmembrane localization of this helix does not seem to be required for the expression of ABCB5β_P-gp linker_ABCB6, either in insect cells or for ATPase activity. This is consistent with the three-dimensional model of ABCB5β/B6 and the conclusion made by Kiss and colleagues, who showed that the disruption of ABCB6 TMD0 leads to mislocalization without impairing either its dimerization or ATPase activity(38). Similar results were obtained for ABCB9 TMD0; its expression was necessary for lysosomal targeting but not for transport function and dimerization(39).

After expression in high-five insect cells, ATPase activity of each heterodimer was investigated. In Figs. 6 and 7, ATPase activity is quantified per mg of membrane protein and is not normalized to the transporter expression. However, when considering the expression levels determined after Coomassie-blue staining quantification, ABCB5β_P-gp linker_ABCB9 displayed the highest basal ATPase activity, followed by ABCB9_P-gp linker_ABCB5β, ABCB6_P-gp linker_ABCB5β, and ABCB5β_P-gp linker_ABCB6. More experiments are needed to evaluate the influence of ABCB5β orientation on protein expression and ATPase activity of the heterodimers. For the homodimers, similar levels of expression and ATPase activities were observed.

We have demonstrated that both heterodimers exhibit basal ATPase activity that was inhibited around 50% by beryllium fluoride, but not by vanadate. Similar results have been reported for ABCB4, ABCG1, ABCC1, and ABCC10, for which vanadate showed little to no inhibitory effect when compared to beryllium fluoride(40-42). At this point, further investigation is needed to identify ABCB5β/B6 and ABCB5β/B9 substrates and determine whether there is any specificity in their substrate spectrum when compared to ABCB5β, ABCB6, and ABCB9 homodimers. It is possible that these heterodimers could have no transport function and only exist to regulate ABCB6 and ABCB9 homodimer formation. Among the mammalian ABC transporters superfamily, previous publications have shown that ABC transporters are able to interact with different partners. For instance, using chimeric dimers, ABCD1 and ABCD2 were able to form functional homodimers, as well as heterodimers, in the peroxisome(43). In HEK293, ABCG4 formed either a homodimer or heterodimer with ABCG1(44). Nevertheless, the physiological relevance of having multiple interaction partners for ABC transporters remains unknown.

The localization of these two heterodimers must be determined and these studies are currently underway. ABCB6 has a conflicting localization in the literature; this transporter has been shown to specifically localize in the outer membrane of the mitochondria(45), and has also been reported to localize in both the outer mitochondrial membrane and the plasma membrane(46). In a more recent study, ABCB6 was reported to have an endolysosomal localization(47). ABCB9 was shown to go through the Golgi, then the endosomes, and to finally reach the lysosomes(48). ABCB5β localization remains elusive but, based on its interacting partners, we can speculate that it is localized in lysosome-related organelles.

In conclusion, this study presents the first evidence for two novel dimers with ATPase activity formed by the heterodimerization of ABCB5β with either ABCB6 or ABCB9 in Mel JuSo and UACC-257 melanoma cell lines. This discovery is a step toward a better understanding of the role of ABCB5β in melanocytes and melanoma.

## Methods

### Cell culture and transfection

The HEK-293T cell line, which is a derivative of human embryonic kidney 293 cells, transformed with the SV40 T-antigen, and the Mel JuSo and UACC-257 melanoma cell lines were cultured in DMEM (Lonza, Basel, Switzerland) supplemented with 10% FBS (GE Healthcare Life Sciences, Issaquah, WA, USA) and 1% penicillin/streptomycin (Gibco, Carlsbad, CA, USA) at 37 °C and 5% CO_2_ in a humidified atmosphere.

For transient transfections, cells were seeded to reach 70% confluence in a six-well plate. Four hours later, transfection with jetPRIME (Polyplus transfection, Illkirch, France) was carried out following the manufacturer’s instructions.

shRNAs constructs for the stable knockdown of human ABCB6 (sc-94721-SH) and ABCB9 (sc-60115-SH) were obtained from Santa Cruz Biotechnology (Dallas, TX, USA). A non-silencing shRNA was used as a control (sc-108060). Reverse transfection was performed in antibiotic-free medium with jetPRIME following the manufacturer’s instructions (Polyplus transfection, Illkirch, France). Seventy-two hours after transfection, the selection of the transfected cells was carried out by the addition of 2 µg/ml of puromycin. After 10 days of selection, ABCB6 and ABCB9 mRNA and protein expression were assessed using RT-qPCR and western blot.

### DNA constructs

For the NanoBRET constructs, ABCB2, ABCB3, ABCB5β, ABCB6, ABCB8, ABCB9, and ABCD1 cDNAs were inserted into the pCDNA3.1 expression vector and subcloned into the NanoLuc pNLF1-N[CMV/Hygro], the NanoLuc pNLF1-C[CMV/Hygro], the HaloTag pHTC HaloTag CMV-neo, and the HaloTag pHTN HaloTag CMV-neo vector purchased from Promega (Madison, WI, USA). Restriction enzymes Not1-EcoR1, Nhe1-EcoR1, Not1-EcoR1, and Nhe1-EcoR1 were used, respectively.

The mCherry-tagged ABCB5 mammalian expression construct (ID: 8488-X02-787), driven by a CMV promoter, was provided by Ledios Biomedical Research (formerly SAIC-Frederick, NCI, MD, USA).

For the heterodimer constructs, ABCB5β with ABCB6 and ABCB5β with ABCB9 coding sequences were fused following the NEBuilder HIFI DNA Assembly kit’s instructions (New England Biolabs, Ipswich, MA, USA). The ABCB5β-ABCB6, ABCB6-ABCB5β, ABCB5β-ABCB9, and ABCB9-ABCB5β sequences were cloned in pDONR201 plasmid. Using the SACII restriction enzyme, the sequence derived from P-gp flexible linker region residues (NEVELENAADESKSEIDALEMSSNDSRSSLIRKRSTRRSVRGSQAQDRKLSTKEALD) 703N-760D, was inserted between each of the half transporters. DNA sequencing was used to confirm both the correct orientation of the linker and absence of other mutations. ABCB5β_P-gp linker_ABCB6, ABCB6_P-gp linker_ABCB5β, ABCB5β_P-gp linker_ABCB9, and the ABCB9_P-gp linker_ABCB5β, ABCB5β, ABCB6 and ABCB9 sequences were transferred into bacmid (pDEST-008), and baculovirus was generated by the Protein Expression Laboratory Cloning and Optimization Group (Frederick National Laboratory for Cancer Research, Frederick, Maryland, USA).

### RT-qPCR

Total RNA was isolated from Mel JuSo and UACC-257 using a NucleoSpin RNA Plus kit following the manufacturer’s instructions (Takarabio, Kusatsu, Japan). Reverse transcription was performed using a High-Capacity cDNA Reverse Transcription kit and 200 ng of total RNA (Thermo Fisher Scientific, Waltham, MA, USA) according to the manufacturer’s instructions. Quantitative PCR was performed using TaqMan master mix (Thermo Fisher Scientific, Waltham, MA, USA) and 18s expression was used to normalize data. The following probes from Thermo Fisher Scientific were used: 18S (Hs03003631_g1), ABCB6 (Hs00180568_m1), and ABCB9 (Hs00608640_m1).

### NanoBRET

Twenty-four hours after the transient transfection of the NanoLuc and HaloTag constructs, the cells were washed with PBS and trypsinized. Cells were counted and diluted in OptiMEM supplemented with 4% FBS to reach a density of 2 × 10^5^ cells/ml. One µl of 0.1 mM HaloTag NanoBRET ligand 618 (Promega, Madison, WI, USA) per ml of cells, or DMSO in the control conditions, was added. One hundred µl of cells with either ligand or DMSO were plated in a 96-well plate (Corning 3917, Thermo Fisher Scientific, Waltham, MA, USA) and were cultured overnight at 37 °C, 5% CO_2_. Twenty-five µl of 100x dilution of NanoBRET NanoGlo substrate (Promega, Madison, MA, USA) was added per well. Plates were shaken for 30 seconds at 400 rpm and then kept in the dark for 30 seconds. Finally, donor emission (447 nm) and acceptor emission (610 nm) were measured at room temperature using a dual filter (SpectraMax i3x, Molecular Devices, San Jose, CA, USA). For all experiments, biological triplicates were performed. All biological triplicates were composed of technical triplicates.

Relative light units (RLU) obtained for donor and acceptor emission were processed as follows to obtain milli Bioluminescence Units (mBU):

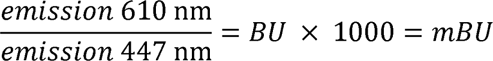

For saturation experiments, percentages of the maximum NanoBRET ratio were plotted against the acceptor to donor ratio; data were fitted using the “one site–total and nonspecific binding” fit in Prism.

### Co-immunoprecipitation

Five mg of total proteins (from either HEK93T, Mel Juso, or UACC-257) in lysis buffer (120 mM NaCl, 50 mM Tris, NP40 0.5%, 5 mM EDTA, 1x protease inhibitor) were incubated on a rotating wheel at 4 °C for 4 hours with 1 µg of antibody: ABCB5 (600-401-A775, Rockland Immunochemicals, Pottstown, PA, USA), ABCB6 (G-10, Santa Cruz Biotechnology, Dallas, TX, USA), ABCB9 (A-8, Santa Cruz Biotechnology, Dallas, TX, USA), IgG Mouse (sc-2025, Santa Cruz Biotechnology, Dallas, TX, USA), and IgG Rabbit (sc-3888, Santa Cruz Biotechnology, Dallas, TX, USA). Simultaneously, 25 µl of magnetic beads, Dynabeads-Protein A (Invitrogen, Waltham, MA, USA), and Dynabeads-Protein G (Invitrogen, Waltham, MA, USA) were added to an Eppendorf and washed twice with 250 µl of lysis buffer using a magnetic rack. Thereafter, protein lysates with the antibody were added to the beads and incubated overnight at 4 °C on a rotating wheel. The following day, the beads were washed three times with 250 µl of cold PBS 0.05% Tween 20. Fifty µl of elution buffer (DTT, Tris, Glycerol, SDS) were added, and samples were heated at 40 °C for 10 minutes. Samples were then analyzed using western blot.

### SDS-PAGE and Western blotting

Samples were loaded on an 8% SDS acrylamide gel, migrated at 110 V, and transferred to a PVDF membrane at 110 V for 1h30. The membrane was blocked for 1 hour in 5% milk diluted in PBS 0.05% Tween 20 and incubated overnight with the following primary antibodies: anti-ABCB5 (600-401-A775, Rockland Immunochemicals, Pottstown, PA, USA), anti-ABCB6 (sc-365930, Santa Cruz Biotechnology, Dallas, TX, USA), anti-ABCB9 (sc-39412, Santa Cruz Biotechnology, Dallas, TX, USA), and anti-mCherry (ab167453, Abcam, Cambridge, UK). Then, membranes were incubated for 1 hour with either Scan Later anti-mouse or anti-rabbit secondary antibodies (Molecular Devices, San Jose, CA, USA); they were then washed with PBS 0.05% Tween 20 and revealed with the SpectraMax i3x (Molecular Devices, San Jose, CA, USA). Alternatively, membranes were incubated for 1 hour with horseradish peroxidase-conjugated goat anti-mouse or anti-rabbit antibodies (Agilent Technologies, Santa Clara, CA, USA), washed with PBS 0.05% Tween, and revealed with enhanced chemiluminescence (PerkinElmer, Waltham, MA, USA).

### *In situ* proximity ligation assay

Cells grown on a coverslip were fixed in 4% paraformaldehyde (PFA) for 20 minutes and permeabilized with PBS + Triton X-100 0.2% for 10 minutes. Then, proximity ligation assay (PLA) was performed in accordance with the Duolink kit (Sigma-Aldrich, Saint Louis, MO, USA). Briefly, the coverslips were incubated with blocking solution for 1 hour at 37 °C in a humidity chamber. Then, they were incubated for 1 hour at room temperature with primary antibody pairs raised against different species (mouse and rabbit) and diluted 1/200: ABCB5β (LS-C169144, LSBio, Seattle, WA, USA), ABCB6 (600-401-945, Rockland Immunochemicals, Pottstown, PA, USA), ABCB8 (ab83194, Abcam, Cambridge, UK), and ABCB9 (PA5-99017, Invitrogen, Waltham, MA, USA). Cells were incubated with PLA probes mouse PLUS and rabbit MINUS for 1 hour at 37 °C. Next, ligation was performed for 30 minutes at 37 °C, followed by amplification for 100 minutes at 37 °C. Finally, the coverslips were mounted on mounting medium with DAPI (Sigma-Aldrich, Saint Louis, MO, USA). The edges were sealed with clear nail polish. Images were obtained using a Leica TCS SP5 confocal microscope (Leica Microsystems, Wetzlar, Germany). DAPI, argon, and 561 red filters were used. The number of red dots per cell was counted using ImageJ software as described in Gomes et al(49). Each condition was obtained in biological triplicates.

### Atomic models

An accurate alignment between the human ABCB5β and mouse ABCB1 sequences was generated based on a multi-sequence alignment that includes various members of the ABCB family transporters. Based on this alignment, the structural model of ABCB5β was constructed manually using the structural model of mouse P-gp (PDB:5KPI)(50) as a template, replacing the residues of mouse ABCB1 with the corresponding residues of ABCB5β in the molecular graphics program Coot(51). The model was subsequently subject to energy minimization using the crystallographic refinement program Refmac(52).

The construction of the heterodimeric ABCB5β-ABCB6 model was initiated using a dimeric cryo-EM model of ABCB6 (PDB:7D7N)(53). A single monomer model of ABCB6 was superposed to the corresponding subunit of the ABCB5β dimer model. Manual adjustments were made to make sure all helices were correctly aligned with the NBDs using the molecular modeling program Coot. The model was then energy minimized using the crystallographic refinement program Refmac. The ABCB5β-ABCB6 model was rendered as ribbon diagram in the program Pymol(54).

The construction of the ABCB5β and ABCB9 heterodimeric model started with the construction of the homology model of ABCB9. The ABCB9 sequence was aligned with that of ABCB6. Mutations were manually introduced into the ABCB6 subunit of the ABCB5β-ABCB6 model based on the sequence alignment in Coot. The resulting ABCB5β-ABCB9 model was then subjected to several cycles of energy minimization.

### Insect cell total membranes preparation

*Trichoplusia ni* (high-five) insect cells were infected at a MOI of 10 with a recombinant baculovirus carrying one of the following sequences: ABCB5β, ABCB6, ABCB9, ABCB5β_P-gp linker_ABCB6, ABCB6_P-gp linker_ABCB5β, ABCB5β_P-gp linker_ABCB9, ABCB9_P-gp linker_ABCB5β. Cells were harvested after 58-72 hours, washed with PBS containing 1% aprotinin, and stored at −80°C. Total membrane vesicles were prepared with hypotonic lysis and differential centrifugation as detailed in (55).

Total membrane vesicles were electrophoresed on a 7% NuPAGE precast gel (Invitrogen, Waltham, MA, USA) for 1h10 at 150 V. Then gels were either stained with Coomassie-blue (15 µg of protein per lane) (Instant blue, abcam, Cambridge, UK) or transferred to nitrocellulose membranes (0.25 µg or 1 µg of protein per lane for the monomeric transporters and heterodimer respectively) and probed with the following primary antibodies overnight at 4°C: anti-ABCB5 (600-401-A775, Rockland Immunochemicals, Pottstown, PA, USA), anti-ABCB1 (C219, Fujirebio, Tokyo, Japan), anti-ABCB6 (sc-365930, Santa Cruz Biotechnology, Dallas, TX, USA), anti-ABCB9 (sc-39412, Santa Cruz Biotechnology, Dallas, TX, USA). Then, nitrocellulose membranes were incubated with horseradish peroxidase-conjugated goat anti-mouse secondary (Santa Cruz Biotechnology, Dallas, TX, USA) for one hour at room temperature and revealed using ChemiDoc (BioRad, Hercules, CA, USA).

### ATPase assay

Total membrane vesicles prepared from high-five insect cells were diluted in ATPase assay buffer (50 mM MES-Tris pH 6.8, 50 mM KCl, 5 mM sodium azide, 1 mM EGTA, 1 mM ouabin, 10 mM MgCl_2_, 2 mM DTT) to reach a final concentration of 10 µg protein/0.1 ml. The reaction was started with the addition of 5 mM ATP and was performed at 37°C for 20 minutes. After 20 minutes, the reaction was terminated with 0.1 ml of 10% SDS. The amount of inorganic phosphate released was measured in both the presence and the absence of Vi (0.3 mM) and BeFx (0.2 mM beryllium sulfate and 2.5 mM sodium fluoride) using a colorimetric method described in (56).

### Statistical analysis

Data were analyzed using GraphPad Prism version 5.04 for Windows with either linear regression or an unpaired Student’s *t*-test. Statistical analysis is described in the figures’ legends. Statistical significance was defined as *p* < 0.05.

## Acknowledgments

We would like to thank Simon Lefèvre for his initial work on co-immunoprecipitations, and Andy Chevigné of the Department of Infection and Immunity, Luxembourg Institute of Health, for the useful conversation regarding NanoBRET experiments. We are grateful to Vincent Van Hée and Marylène Focant of Promega for their support in the optimization and troubleshooting of the NanoBRET assay. We would also like to thank Marielle Boonen (Intracellular Trafficking Biology, NARILIS, UNamur) for her critical reading of the manuscript. P.S., D.X., S.V.A, and M.M.G were supported by the Intramural Program of the National Institutes of Health, National Cancer Institute, Center for Cancer Research.

## Author contributions

L.G. designed and performed the majority of the experiments and analyzed the data. L.D. prepared the NanoBRET constructs. L.D. and L.S. contributed to the NanoBRET assay optimizations. M.F. performed co-IPs using the mCherry-tagged ABCB5β construct. P.S. assisted with the design and preparation of membrane vesicles and ATPase assays with heterodimeric chimeras as well as manuscript editing. D.X. did the three-dimensional models. L.G. and J.P.G. wrote and edited the manuscript. M.M.G. and S.V.A. provided overall direction and helped edit the manuscript. J.P.G. supervised the project and contributed to the design of the experiments and the analysis of the data.

## Conflict of interest

The authors declare that they have no conflicts of interest with the contents of this article.

## Notes

### Competing Interest Statement

The authors have declared no competing interest.

### Summary of Updates

Author list and results presentation updated

