## Supplemental information for "Identification of two novel heterodimeric ABC transporters in melanoma: ABCB5β/B6 and ABCB5β/B9"

### Supporting information

A.

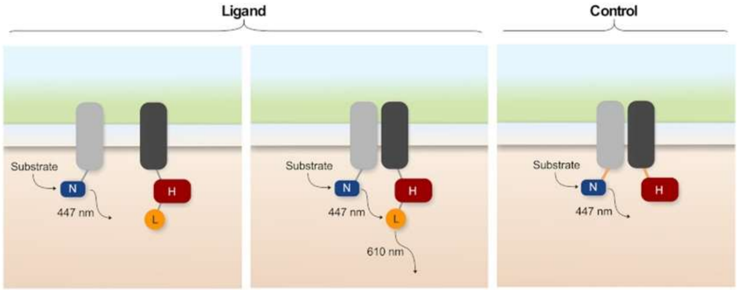


B. **Positive control**

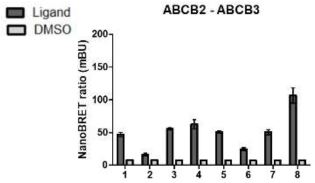


| | ABCB3 | Mean (mBU) |
|---|---|---|
| ABCB2 | ABCB3 NanoLuc C ter – ABCB2 HaloTag C ter | 106.65 |

**Negative controls**

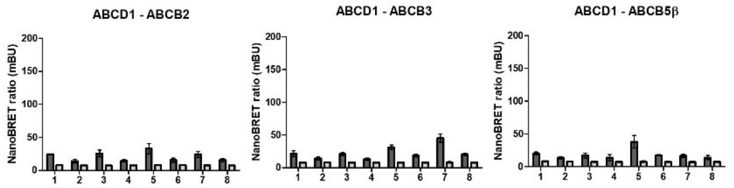


| | ABCD1 | Mean (mBU) |
|---|---|---|
| ABCB2 | ABCB2 NanoLuc N ter – ABCD1 HaloTag N ter | 25.14 |
| ABCB3 | ABCB3 NanoLuc C ter – ABCD1 HaloTag N ter | 37.11 |
| ABCB5β | ABCB5β NanoLuc N ter – ABCD1 HaloTag N ter | 30.30 |

**ABCB5β heterodimerization investigation**

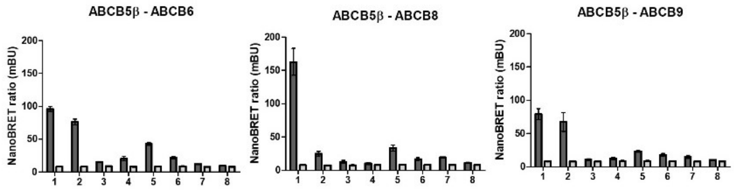


| | ABCB5β | Mean (mBU) |
|---|---|---|
| ABCB6 | ABCB5β NanoLuc N ter – ABCB6 HaloTag N ter | 87.91 |
| ABCB8 | ABCB5β NanoLuc N ter – ABCB8 HaloTag N ter | 155.10 |
| ABCB9 | ABCB5β NanoLuc N ter – ABCB9 HaloTag N ter | 79.16 |

**Fig. S1: Schematic representation of the NanoBRET technique and preliminary NanoBRET ratio**. **A** NanoBRET consists of an engineered luciferase, NanoLuc (N), fused to the transporter of interest. The HaloTag protein (H) is fused to the potential interacting partner. Following substrate addition, NanoLuc emits at 447 nm. If the HaloTag ligand (L), covalently bound to the HaloTag protein, is in close proximity (less than 10 nm), it will be excited by NanoLuc emission and will emit at 610 nm. The ratio between the 447 nm and 610 nm signals allows distinction between interacting (i.e., high ratio) and noninteracting proteins (i.e., low ratio, close to the technical negative control). The same experiment without the ligand serves as the technical negative control. **B** NanoBRET ratios of the positive control, negative controls, and investigation of putative ABCB5β heterodimerization. ABC transporters were cloned in either the NanoLuc (N) or HaloTag (H) vector in either the N or C terminus, allowing for eight possible combinations per transporter's pairs. Data are represented as mean ± SD. The order of the combination on each graph is as follows, with ABC 1 being the first ABC transporter mentioned in the graph title and ABC 2 being the second: 1) ABC 1 NanoLuc N terminus – ABC 2 HaloTag N terminus, 2) ABC 1 NanoLuc N terminus – ABC 2 HaloTag C terminus, 3) ABC 1 NanoLuc C terminus – ABC 2 HaloTag N terminus, 4) ABC 1 NanoLuc C terminus – ABC 2 HaloTag C terminus, 5) ABC 2 NanoLuc N terminus – ABC 1 HaloTag N terminus, 6) ABC 2 NanoLuc N terminus – ABC 2 HaloTag C terminus, 7) ABC 2 NanoLuc C terminus – ABC 1 HaloTag N terminus, 8) ABC 2 NanoLuc C terminus – ABC 1 HaloTag C terminus. One µg of NanoLuc and HaloTag constructs was transfected in HEK-293T. Preferred tag orientations selected for further testing in dilution and saturation assays are highlighted in the tables, along with their mean NanoBRET ratios.

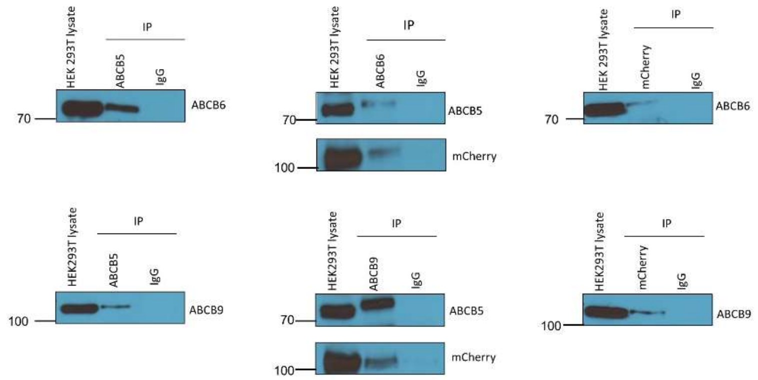


**Fig. S2: Co-immunoprecipitation of ABCB5β-ABCB6 and ABCB5β-ABCB9 using HEK-293T transfected with the mCherry-tagged ABCB5 construct**. Cell lysates were immunoprecipitated (IP) using an anti-ABCB5, anti-ABCB6, anti-ABCB9, or anti-mCherry antibody. The precipitated proteins were visualized by SDS-PAGE using the corresponding antibody indicated on the right side of each blot. Fifty µg of HEK293T whole cell lysate and the total volume obtained after IP were loaded on the gel for the lysate or IP conditions, respectively. An isotype control was performed to determine the specificity of the signal obtained in the Western blots.

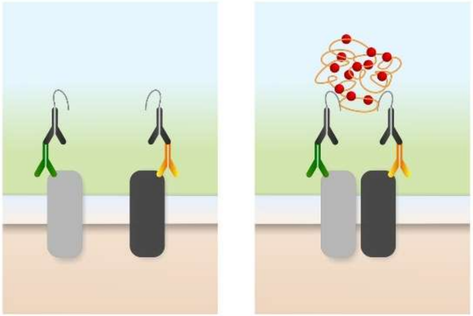

**Fig. S3: Schematic representation of the proximity ligation assay**. The proximity ligation assay consists of the incubation with primary antibody from two different species specific to the target proteins. Then, samples were incubated with secondary antibodies coupled with PLA probes and raised against the desired species. If they are in close vicinity, a connector oligonucleotide joins the PLA probes. This results in a closed circular DNA template, which is amplified using a DNA polymerase. Finally, a detection oligonucleotide coupled to a fluorochrome hybridizes to repeating sequences in the amplicon. The PLA signal was detected using fluorescence microscopy as discrete dots, which illustrate the interaction between the proteins of interest.

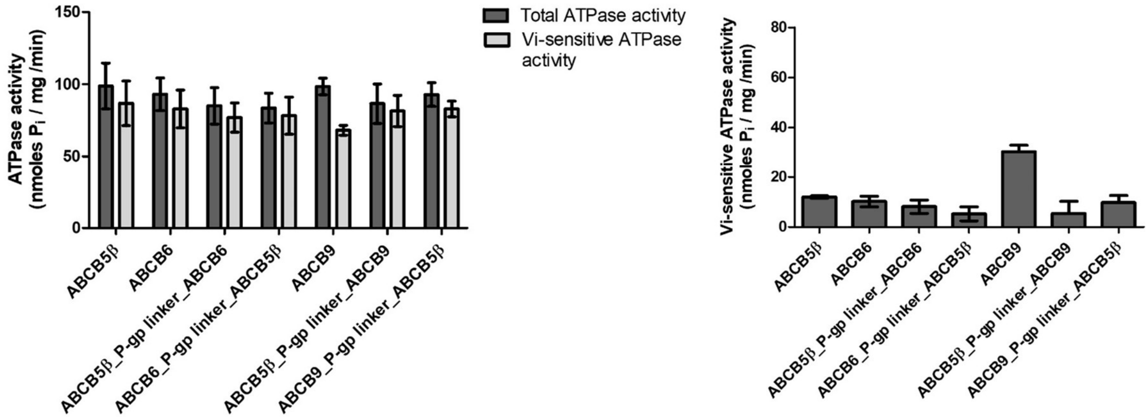


**Fig. S4: Vi-sensitive ATPase activity of homo- and heterodimeric transporters**. ABCB5β, ABCB6, ABCB5β_P-gp linker_ABCB6, ABCB6_P-gp linker_ABCB5β, ABCB9, ABCB5β_P-gp linker_ABCB9, and ABCB9_P-gp linker_ABCB5β ATPase activity was measured in the presence of Vi as described in the method section. On the left, data are represented as ATPase activity in absence of Vi (total ATPase activity) and presence of Vi (Vi-sensitive ATPase activity). On the right, data are represented as the difference between Vi ATPase activity and total ATPase activity (n = 3).

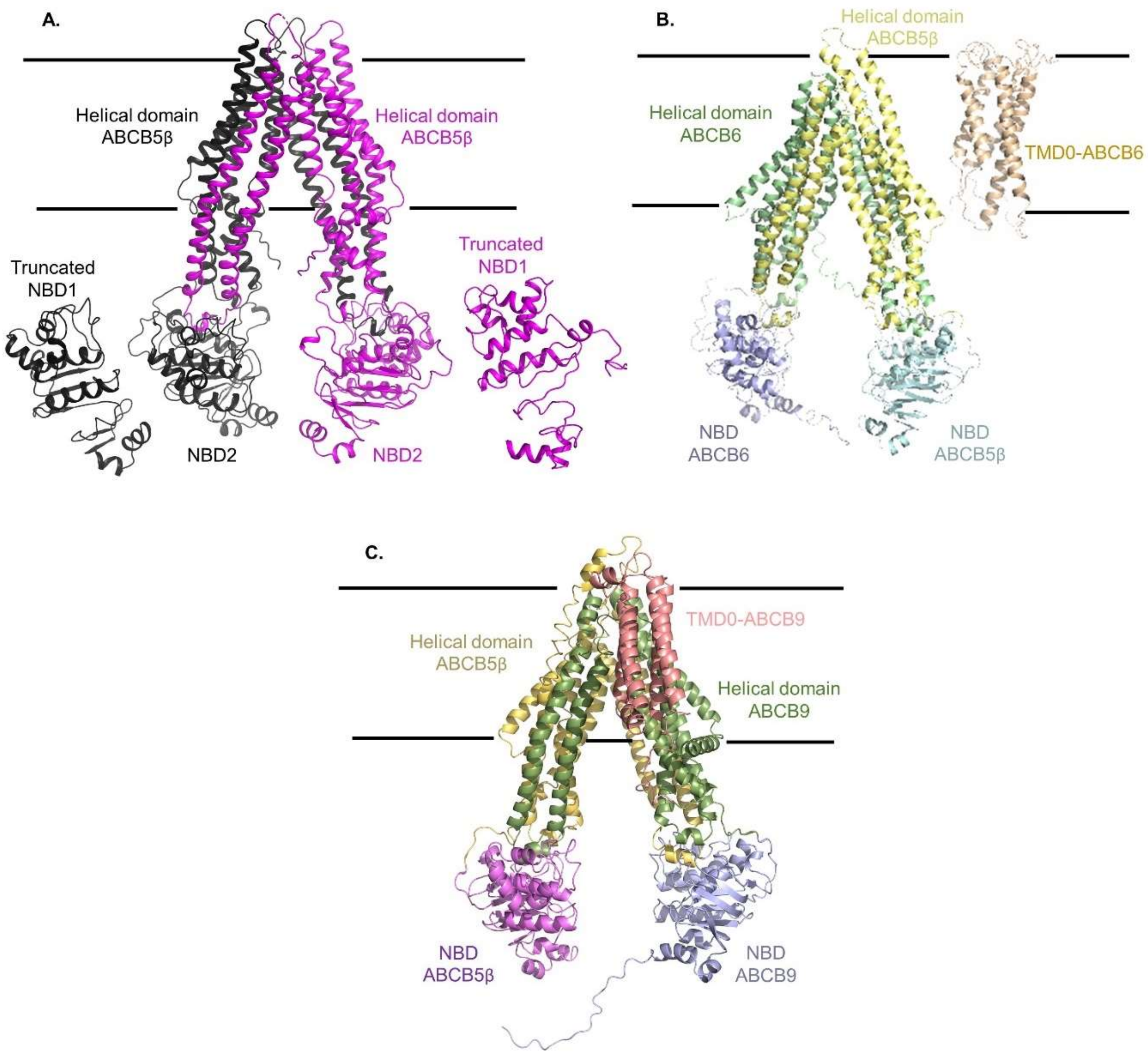


**Fig. S5: Model structures of ABCB5β/B5β, ABCB5β/B6 and ABCB5β/B9.** The model structures of ABCB5β/B5β, ABCB5β/B6, and ABCB5β/B9 in an inward-open conformation are illustrated as cartoon representation. The two horizontal lines represent the boundary of the membrane bilayer. (**A**) Structure of the ABCB5β-ABCB5β homodimer. The first ABCB5β monomer is in black and the second is in purple. Truncated NBD1 and NBD2 for each monomer are labelled. (**B**) Structure of the ABCB5β-ABCB6 heterodimer. The helical domain of the ABCB5 is yellow and the NBD cyan. The model of ABCB6 consists of three domains: a TMD0 shown in coral, a helical domain in green, and an NBD in purple. (**C**) Structure of the ABCB5β-ABCB9 heterodimer. The helical domain of ABCB5 is yellow and its NBD is magenta. The model of ABCB9 is comprised of three domains: a TMD0 in brown, a helical domain in green, and an NBD domain in purple.
